# Alternative activation of macrophages is accompanied by chromatin remodeling and short-term dampening of macrophage secondary response

**DOI:** 10.1101/327023

**Authors:** Mei San Tang, Emily R. Miraldi, Natasha M. Girgis, Richard A. Bonneau, P’ng Loke

## Abstract

Interleukin-4 (IL-4) activates macrophages to adopt a distinct phenotype associated with clearance of helminth infections and tissue repair. Here, we describe changes in the accessible chromatin landscape following IL-4 stimulation of terminally differentiated mouse peritoneal macrophages. This chromatin remodeling process occurs in both tissue resident and monocyte-derived macrophages, but the regions gaining accessibility post-stimulation are macrophage-specific. PU.1 motif is similarly associated with tissue resident and monocyte-derived IL-4 induced regions, but has macrophage-specific DNA shape and predicted co-factors. In addition, IL-4 stimulation leads to short-term dampening of macrophage secondary response. However, the degree of dampening differs between macrophages derived from different genetic backgrounds. Together, these results lead us to propose that DNA sequence variations can alter parts of the accessible chromatin landscape and differences in secondary responses due to host genetics can contribute to phenotypic variations in immune responses.

## Introduction

Macrophage activation is a process by which macrophages transition from a resting state to adopt different phenotypes, in response to specific external stimuli that can either be danger signals or homeostatic and metabolic signals (1). The macrophage activation process is accompanied by changes in transcriptional activities and histone modifications genome-wide, orchestrated by combinatorial actions of different transcription factors (TFs) that include lineage-determining TFs (such as PU.1) and stimulus-dependent TFs (such as the STAT and IRF proteins) (1-6). However, such molecular events have almost exclusively been described for bone marrow derived macrophages (BMDMs) in response to toll-like receptor (TLR) signaling, which is often used as a reductionist model to mimic type 1 immune response to acute infections that gives rise to classically activated macrophages.

In contrast, alternatively activated macrophages (AAMs) induced by type 2 cytokines, such as interleukin-4 (IL-4) and IL-13, adopt a distinct phenotype that can promote helminth expulsion and limit tissue damage during helminth infection (7-10). We have previously demonstrated that macrophages of tissue resident and monocytic origins are phenotypically different following IL-4 stimulation (11, 12). Here, we expand on these macrophage-specific differences by characterizing changes in accessible chromatin landscape following IL-4 stimulation of these different types of macrophages. The effects of IL-4 on BMDMs have also been documented, particularly highlighting the reduced response to interferon gamma after IL-4 stimulation that is mediated by the action of TFs such as the PPARγ:RXR heterodimer and STAT6 (13-15). However, the effects of IL-4 on the chromatin of macrophages from different cellular lineages *in vivo* have yet to be carefully investigated.

*In vivo*, chromatin accessibility changes have mostly been associated with the cellular differentiation process, but we find that IL-4 stimulation alone can give rise to new accessible regions in terminally differentiated peritoneal macrophages. These IL-4 induced regions are macrophage-specific, with differences in TF motifs. In addition, IL-4 stimulation *in vivo* leads to a short-term dampening of macrophage response to a repeat IL-4 stimulation. While there have been considerable number of studies that described the opposing effects of different cytokines (16, 17), none has so far demonstrated that a single cytokine can dampen its own response upon repeated stimulation. Notably, this dampening in response to the second IL-4 stimulation occurs to different degrees in C57BL/6 and BALB/c mouse strains. These differences may be important for the outcome of helminth infections, since during *Litomosoides sigmodontis* infection the more susceptible BALB/c mice undergo less tissue resident macrophage expansion and more monocyte infiltration than the resistant C57BL/6 strain (10). Finally, because our knowledge of macrophage biology is largely based on the C57BL/6 mouse strain, our study highlights the need to account for genetic diversity in mouse immune models.

## Results

### IL-4 stimulation leads to remodeling of open chromatin landscape in peritoneal macrophages

To examine chromatin remodeling on different types of tissue macrophages, we injected recombinant IL-4–antibody complex (IL-4c) into the peritoneal cavity of C57BL/6 mice to induce accumulation of alternatively activated F4/80^hi^CD206^-^ macrophages of embryonic origin (AAM^res^) and compared these with F4/80^int^CD206^+^ macrophages derived from Ly6C^hi^ inflammatory blood monocytes (AAM^mono^) in mice injected with IL-4c and thioglycollate (12). We then used ATAC-seq (18) to profile the open chromatin landscape of these macrophages, in comparison to non-stimulated F4/80^hi^CD206^-^ macrophages of naïve mice and F4/80^int^CD206^+^ macrophages from thioglycollate-treated mice (12).

The overall differences in accessible chromatin landscape (a total of 61,713 open chromatin regions) could be attributed mainly to the type of macrophage (27% of total variance), but alternative activation by IL-4 also altered the accessible chromatin profiles (Figures 1A, B). *Arg1* and *Ucp1*, which are known to be IL-4 inducible (12), had constitutively accessible chromatin regions, whereas *Retnla*, another IL-4 inducible gene, had chromatin regions that gained accessibility in response to IL-4 (Figure 1B). This IL-4 induced chromatin remodeling process can be cell-type-specific at certain regions (e.g. regions adjacent to the loci of *Tgfb2, Ccl2*) (Figure 1B). Of the 61,713 total accessible regions, we identified 1572 regions induced by IL-4 for AAM^res^ and 1462 regions for AAM^mono^ (Figure S1A). IL-4-dependent regions also had the largest contribution to the differences in open chromatin profiles between non-stimulated and IL-4 stimulated macrophages (Figure 1C).

**Figure 1:**
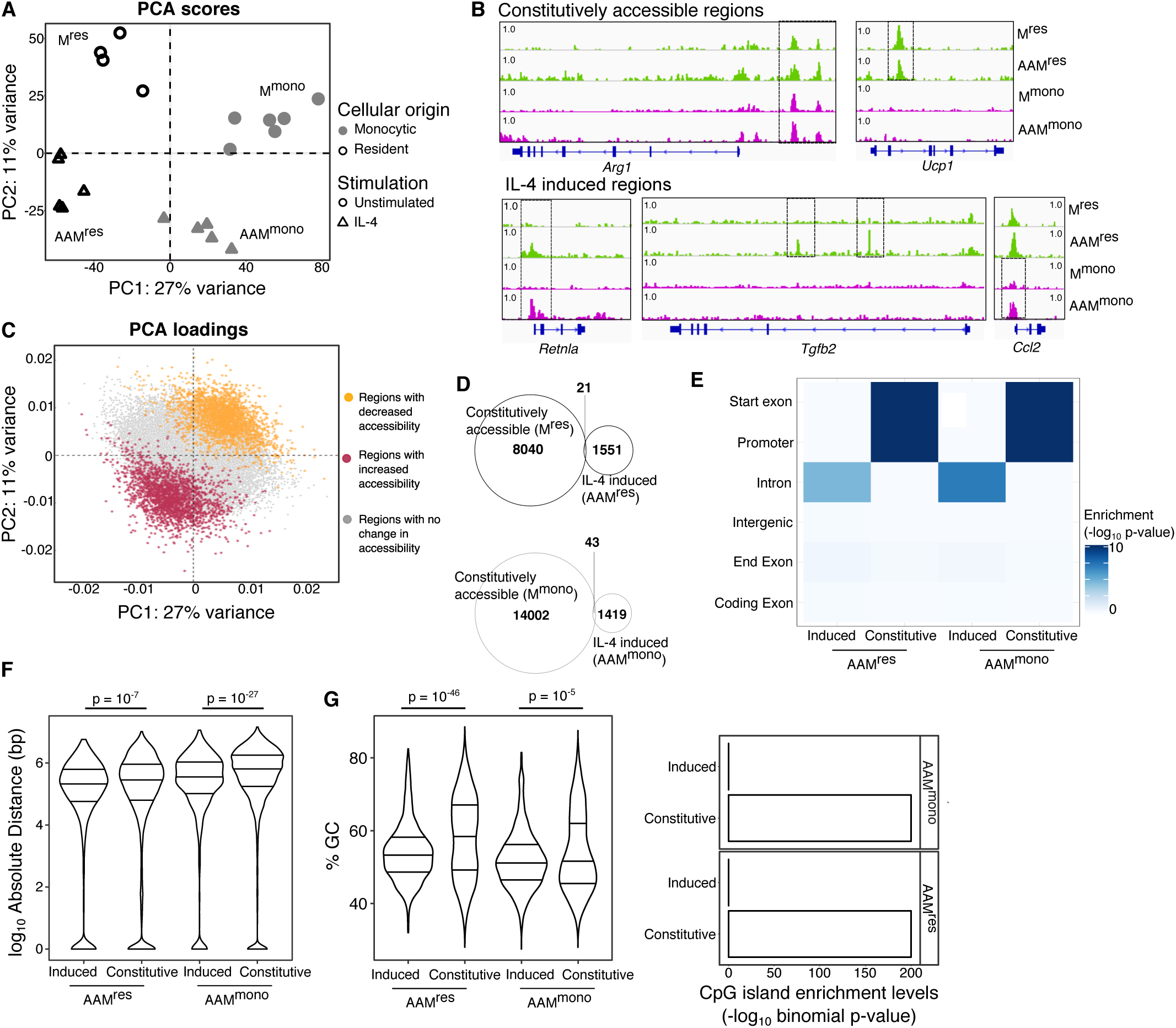
IL-4 stimulation leads to remodeling of open chromatin landscape in peritoneal macrophages. (A) PCA scores of individual ATAC-seq samples. PCA was performed using rlog-transformed ATAC-seq read counts of 30,856 regions with high variance (only regions with variance inter-quartile range > 0.5 were retained). N = 4-6 mice per macrophage population. Data points represent independent biological replicates. (B) Genome browser views of representative (boxed) constitutively accessible and IL-4 induced regions. (C) The contributions of individual accessible regions to PCs 1 and 2 are represented in the PCA loadings plot. Each data point is color-coded based on the direction of its IL-4 dependency. Hence, IL-4 induced regions (red) are highly associated with IL-4 stimulated macrophages, while IL-4 repressed regions (yellow) are highly associated with non-stimulated macrophages. We compared (D) enrichment levels for different types of genomic elements, (E) distance from a closest IL-4-induced gene and (F) G/C content between constitutively accessible and IL-4-induced regions in AAM^mono^ and AAM^res^, respectively. G/C content information is represented in two different ways – percentage of G/C bases in an accessible region (F, left panel) and CpG island enrichment for a given group of accessible regions (F, right panel). Number of IL-4 induced regions = 1572 in AAM^res^ and 1462 in AAM^mono^; number of constitutively accessible regions = 8061 in AAM^res^ and 14045 in AAM^mono^. Enrichment p-values are from binomial test while two-class comparison p-values are from two-sided Mann-Whitney test.

The IL-4 induced regions almost all (99% in AAM^res^ and 97% in AAM^mono^) gained accessibility from undetectable levels at baseline (Figure 1D). We made comparisons of several sequence characteristics between constitutively accessible and IL-4 induced regions (Figure S1B). IL-4 induced regions were more likely to reside in non-coding intronic regions – in tissue-resident macrophages, 781 of 1572 IL-4 induced peaks were intronic, as compared to 2199 of 8061 constitutively accessible regions (binomial enrichment, p = 2.5 × 10^−5^), while in monocyte-derived macrophages, 761 of 1462 IL-4 induced peaks were intronic, as compared to 5282 of 14045 constitutively accessible regions (binomial enrichment, p = 8.5 × 10^−4^) (Figure 1E). In the monocyte-derived macrophages, IL-4 induced regions were overall closer to IL-4 induced genes (two-sided Mann-Whitney test, p = 3.4 × 10^−25^ in AAM^mono^) (Figure 1F). In tissue-resident macrophages, IL-4 induced regions contained lower GC content (two-sided Mann-Whitney test, p = 8.6 × 10^−46^ in AAM^res^) (Figure 1G, left) and were also less likely to overlap with a CpG island (Figure 1G, right). Hence, IL-4 stimulation can lead to reorganization of the chromatin landscape in terminally differentiated peritoneal macrophages, giving rise to newly accessible regions that have distinct sequence properties when compared to constitutively accessible regions.

### IL-4 induced regions are associated with PU.1, KLF and AP-1 motifs

Even though both AAM^mono^ and AAM^res^ received the same stimulation within the peritoneal tissue environment, the regions that were remodeled by IL-4 were largely macrophage-specific (Figure 2A). Of all the 2855 IL-4 induced regions, only 179 regions (6% of total IL-4 induced regions) were shared between both AAM^mono^ and AAM^res^ (Figure S1C). While the IL-4 induced regions from AAM^mono^ and AAM^res^ were largely distinct, the DNA motifs discovered from these distinct regions were grouped into similar families of TFs, which included PU.1, KLF and the AP-1 family of motifs (Figure 2B). However, AAM^mono^ had significantly higher number of accessible regions with the AP-1 motif (two-sided Fisher’s exact test, p = 9.5 × 10^−10^), while AAM^res^ had significantly higher number of accessible regions with the KLF motif (two-sided Fisher’s exact test, p = 9.2 × 10^−8^), suggesting the use of different TFs by the two macrophage types during chromatin remodeling upon IL-4 induced alternative activation (Figure 2C). Such differences were not observed with the PU.1 motifs. We next examined the expression levels of these TF families (KLF vs. AP-1) to determine if specific members within each family could be differentially expressed between AAM^res^ and AAM^mono^. 20 TFs of KLF and AP-1 families were highly expressed in peritoneal macrophages (Figure S2A). These TFs almost all demonstrated lineage-specific expression, both at baseline (Clusters 1 and 2) and with IL-4 stimulation (Clusters 3 and 4) (Figure 2D). These results indicate that while KLF and AP-1 family of TFs may have lineage-specific functions, PU-1 is likely important for both macrophage lineages.

**Figure 2:**
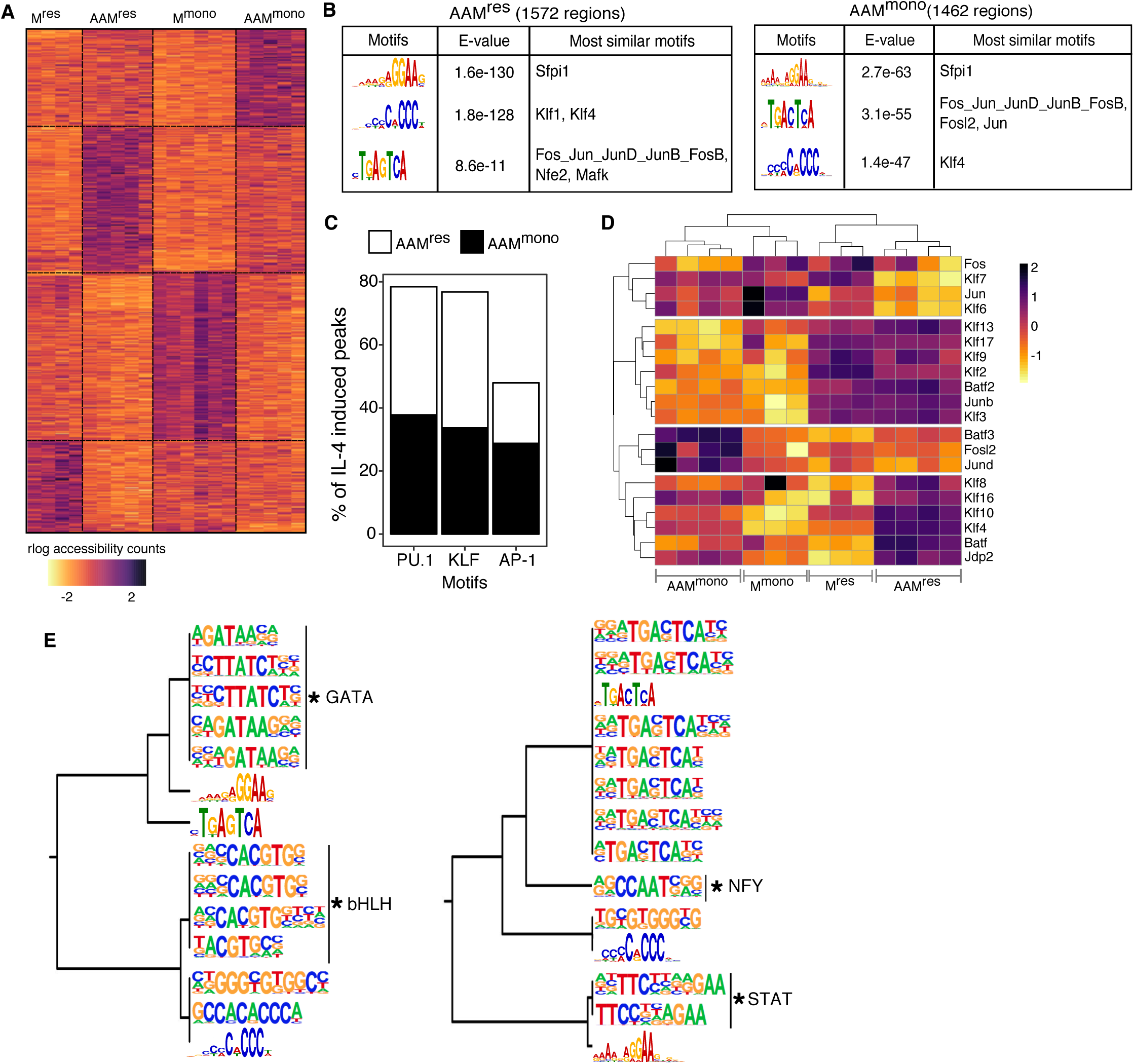
IL-4 induced regions are associated with PU.1, KLF and AP-1 motifs. (A) Heatmap visualizing the macrophage-specific IL-4 dependent regions. Each row represents one of the 2855 IL-4 dependent regions and each column a unique sample. Values are rlog-transformed, batch-subtracted read counts, scaled using a z-score transformation for each region. (B) Motifs discovered *de novo* from IL-4 induced regions in AAM^res^ and AAM^mono^. (C) Frequency of IL-4 induced peaks delineated by the presence of *de novo* PU.1, KLF and AP-1 motifs. (D) 20 highly expressed TF genes related to the *de novo* motifs discovered from the IL-4 induced regions. Values are log_2_ intensity values of microarrays (12). (E) Clustering analysis of *de novo* motifs and macrophage-specific motifs identified using an over-representation approach, from the IL-4 induced regions of AAM^res^ (left) and AAM^mono^ (right). Asterisks indicate macrophage-specific motifs uniquely identified via the over-representation approach. Only macrophage-specific motifs with log_2_ p-value < −15 are included in this visualization and the complete list of motifs identified by over-representation are included as Data S1.

We next used an over-representation approach to identify additional motifs enriched in the IL-4 induced regions of AAM^res^ or AAM^mono^ (Data S1). Since there were some overlaps between the motifs discovered by the *de novo* discovery method and over-representation method, we combined the two sets of motifs for a clustering analysis and merged motifs that were redundant. Using this approach, we identified macrophage-specific motifs beyond those from *de novo* motif discovery (Figure 2E). The GATA motifs and basic helix-loop-helix (bHLH) motifs were specific to IL-4 induced regions of AAM^res^. TFs with these binding motifs have been implicated to be important in proliferation of tissue resident macrophages (19, 20). In contrast, the NFY and STAT motifs were specific to IL-4 induced regions of AAM^mono^. These macrophage-specific motifs were only detected in 15-21% of IL-4 induced regions (236 of 1572 IL-4 induced peaks in AAM^res^ and 317 of 1462 IL-4 induced peaks in AAM^mono^), while the PU.1, KLF and AP-1 motifs discovered by the *de novo* method were present in approximately 75% of IL-4 induced peaks in AAM^res^ and AAM^mono^. Therefore, while there are specific TF motifs enriched in IL-4 induced regions of AAM^res^ (GATA and bHLH) and AAM^mono^ (NFY and STAT), the majority of IL-4 induced regions are enriched for a common set of TF motifs.

### PU.1 motifs in AAM^res^ and AAM^mono^ are associated with macrophage-specific sequence features

PU.1 motif was the most frequently found motif from the IL-4 induced regions across both AAM^res^ (639 of 1572 IL-4 induced regions) and AAM^mono^ (552 of 1462 IL-4 induced regions) (Figure 2C). We focused on these predicted PU.1 binding sites and further characterized their local sequence features in both types of macrophages. We first quantified the accuracy of PU.1 motif prediction with actual PU.1 binding, by comparing our predicted PU.1 motifs from thioglycollate-elicited macrophages with PU.1 ChIP-seq in the same cell type (5, 21). 78% of the PU.1 motif sites predicted from thioglycollate-induced macrophages in our study (4,282 of total 5,492 predicted PU.1 motif sites) overlapped with a PU.1 binding site defined by ChIP-seq (Figure S2B).

We next characterized the PU.1 motifs discovered from the IL-4 induced regions of AAM^res^ and AAM^mono^. Motif score is commonly used as a proxy of TF-DNA binding affinity. When compared to PU.1 motifs from AAM^res^, PU.1 motifs from AAM^mono^ had significantly lower motif scores and also demonstrated greater variability in their values (Figure 3A). To identify potential co-factors that could bind in collaboration with PU.1 and contribute to macrophage-specific PU.1 accessibility, we performed motif scanning using sequences from PU.1 motifs ±25bp flanking sequences. We identified TF motifs that were specific for AAM^res^ vs. AAM^mono^ in these PU.1 regions (Figure 3B, Data S2). These predicted macrophage-specific co-factors are largely from different families. We observed a greater diversity in TF families enriched around the IL-4 induced PU.1 motifs of AAM^mono^. These results indicate that PU.1 may function cooperatively with different co-factors to bind different genomic regions depending on the macrophage lineage.

**Figure 3:**
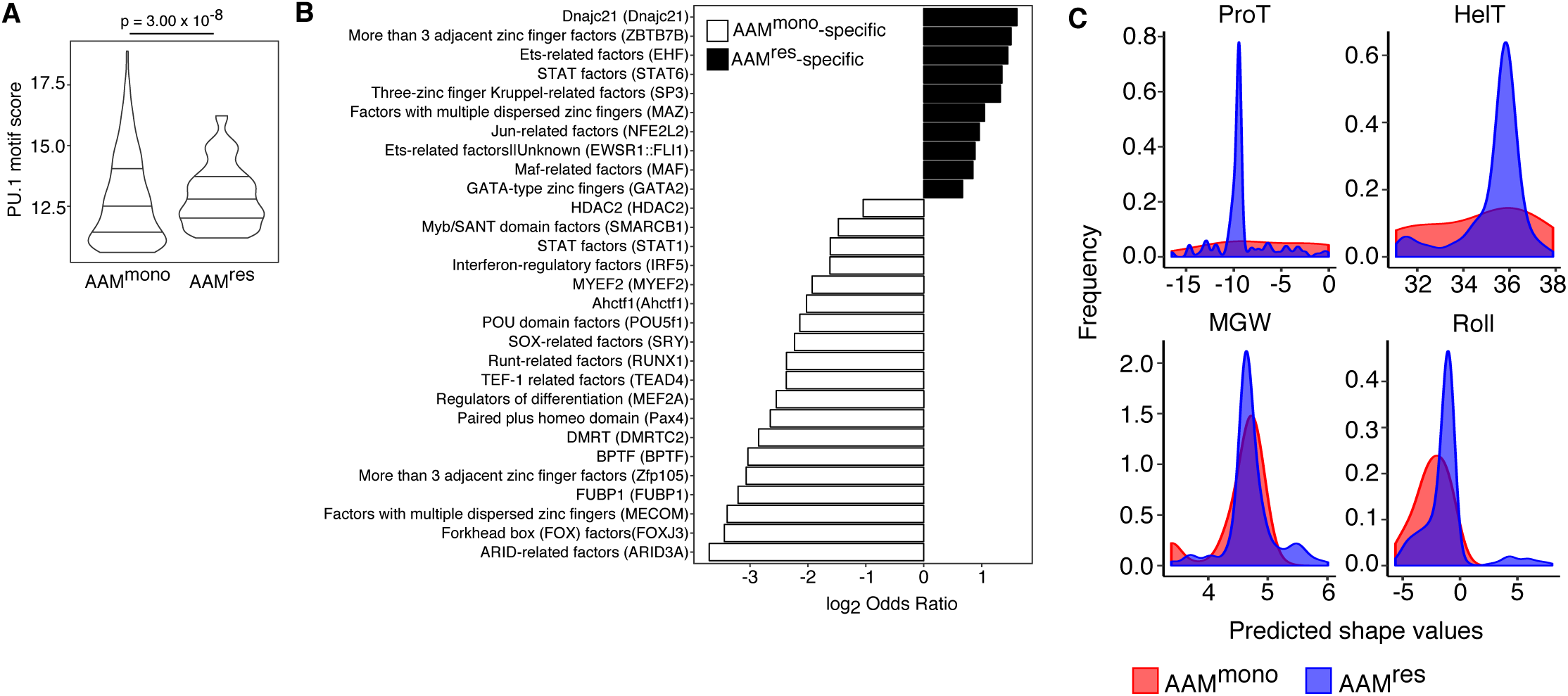
PU.1 motifs in AAM^res^ and AAM^mono^ are associated with macrophage-specific sequence features. (A) Comparison of PU.1 motif scores derived by FIMO in IL-4 induced regions of AAM^res^ vs. AAM^mono^, with horizontal lines in the violin plots representing values at 25^th^, 50^th^ and 75^th^ percentiles. P-value is from a two-sided Mann-Whitney test. Number of IL-4 induced regions = 1572 in AAM^res^ and 1462 in AAM^mono^. (B) Macrophage-specific TF motifs found within IL-4 induced PU.1 motifs ± 25bp regions, represented using log2 odds ratio values (two-sided Fisher’s test, adjusted p-values < 0.1). Motifs are summarized as TF families and the specific TF with the maximum absolute log_2_ odds ratio value is stated in parenthesis. Where TF family annotation was not available, the log_2_ odds ratio of the specific TF itself is used. (C) Predicted DNA shape at the 8^th^ base pair of PU.1 motif of AAM^res^ and AAM^mono^. Frequency distributions are represented by smoothed kernel density estimates. Complete comparisons at all nucleotides are presented in Data S3. ProT = propeller twist, HelT = helical twist, MGW = minor groove width.

Since DNA shape has been shown to be a predictor of TF binding pattern (22, 23), we computationally predicted four DNA shape configurations (minor groove width, propeller twist, helical twist and roll) at the IL-4 induced PU.1 regions of AAM^res^ and AAM^mono^ (24). The PU.1 motifs of AAM^res^ have a more conserved DNA configuration, while PU.1 motifs of AAM^mono^ demonstrated greater variability in DNA configuration. These differences were most pronounced in the propeller and helical twist configurations of the 8^th^ base pair in the PU.1 motif (Figure 3C, Data S3). In summary, while the PU.1 motif is the most enriched motif in the IL-4 induced regions for both AAM^res^ and AAM^mono^, sequence characteristics in the PU.1 motifs differ between the two types of peritoneal macrophages, suggesting PU.1 might cooperate with distinct co-regulators in these different lineages.

### AAMs from C57BL/6 and BALB/c mice are functionally distinct

We next characterized the lineage-specific response to IL-4 stimulation on a different genetic background, by comparing the transcriptional profiles of C57BL/6 and BALB/c AAMs. Most of the differences in transcriptional profiles were driven by macrophage types (45% of total variance), although strain differences also contributed to the considerable variation in transcriptional profiles (18% of total variance) (Figure 4A). Consistent with this finding, most of the macrophage-specific functions were conserved across mouse strains and not affected by genetic differences (Figure 4B, left panel). In contrast, functional differences secondary to genetics were largely specific to the different types of macrophages (Figure 4B, right panel).

**Figure 4:**
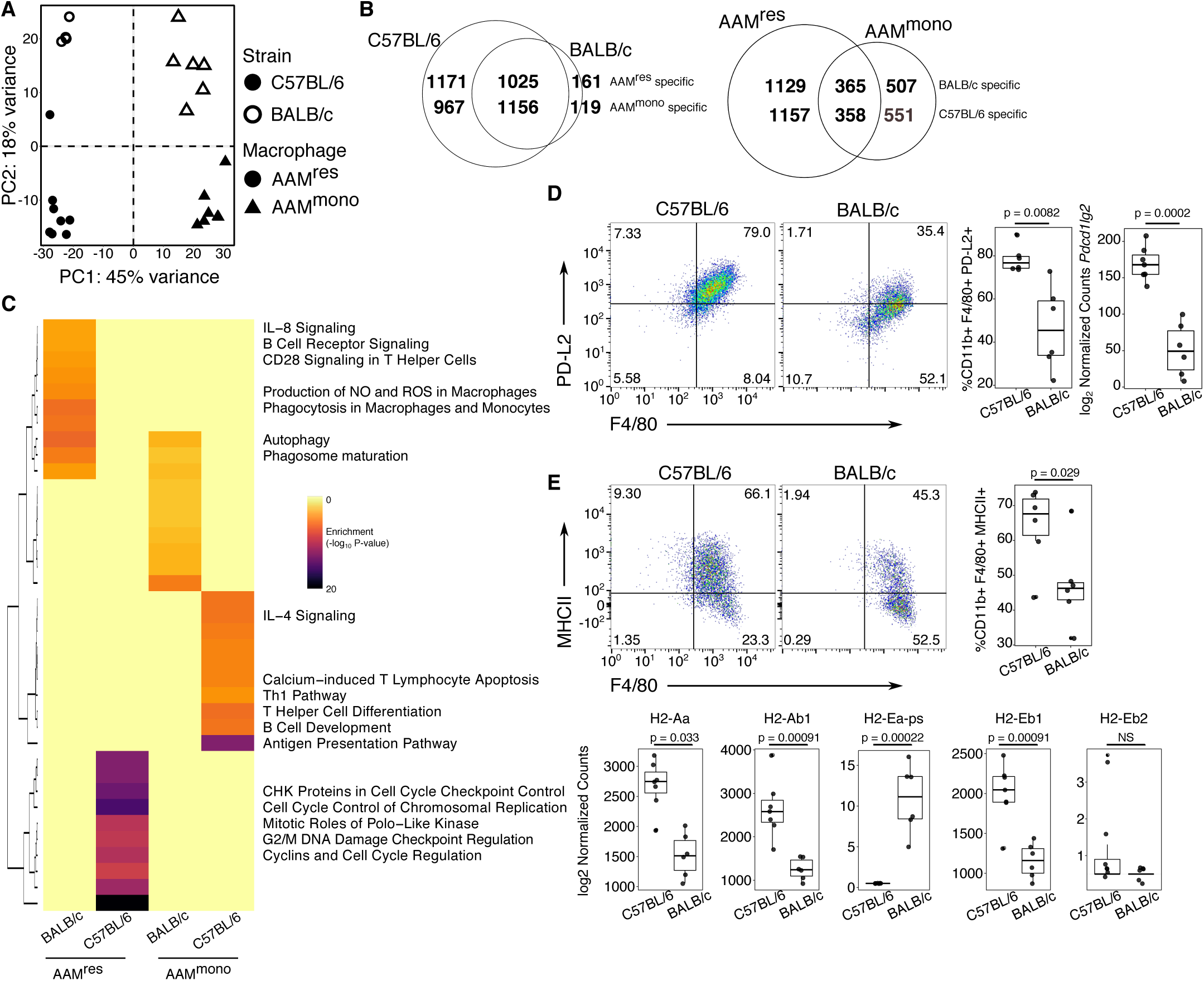
AAMs from C57BL/6 and BALB/c are functionally distinct. (A) PCA of 7431 genes with high variance (only genes with variance inter-quartile range > 0.5 were retained). Data points represent independent biological replicates. (B) Venn diagrams indicating the number of genes that were commonly and uniquely identified as significantly differential (FDR < 0.1) in different comparisons – (left) macrophage-specific genes in C57BL/6 and BALB/c AAMs; (right) strain-specific genes in AAM^res^ and AAM^mono^. (C) Enrichment values from Ingenuity Pathway Analysis visualized as –log_10_ P-value for the four different groups of genes – (1) BALB/c specific in AAM^res^, (2) C57BL/6 specific in AAM^res^, (3) BALB/c specific in AAM^mono^ and (4) C57BL/6 in AAM^mono^. Only the top 10 pathways (as defined by enrichment p-values) are included in this visualization. Specific pathways are highlighted for clarity. The complete lists of enriched pathways are provided as Supplemental Materials Data S2. (D) Representative flow cytometric analysis of F4/80 and PD-L2 surface expressions in AAM^mono^ of C57BL/6 and BALB/c mice. Boxplots show frequency of CD11b+ F4/80+ PD-L2+ singlet, live cells. P-value is based on a two-sided unpaired t-test. (E) Expression of the *Pdcd1lg2* gene in AAM^mono^ of C57BL/6 vs. BALB/c mice, represented by size-factor normalized read counts. P-value is from DESeq2 and adjusted by Benjamini-Hochberg procedure. (F) Representative flow cytometric analysis of F4/80 and MHCII surface expressions in AAM^mono^ of C57BL/6 and BALB/c mice. Boxplots show frequency of CD11b+ F4/80+ MHCII+ singlet, live cells. P-value is based on a two-sided unpaired t-test. (G) Expression values of all MHCII genes in AAM^mono^ of C57BL/6 vs. BALB/c mice, represented by size-factor normalized read counts. P-values are from DESeq2 and adjusted by Benjamini-Hochberg procedure. Hinges of all boxplots correspond to values of the 25th, 50th and 75th percentiles, while boxplot whiskers extend to no more than 1.5 × inter-quartile range, beyond which the outlier data points will be plotted individually. Transcriptional profiling analysis: N = 8 AAM^res^ (C57BL/6), 4 AAM^res^ (BALB/c), 7 AAM^mono^ (C57BL/6), 6 AAM^mono^ (BALB/c). Flow cytometric analysis: N = 6 AAM^mono^ (C57BL/6), 6 AAM^mono^ (BALB/c).

We next examined the strain-specific functional differences in AAM^res^ and AAM^mono^, respectively (Figure 4C, Data S4). BALB/c AAM^res^ expressed lower levels of cell-cycle-related genes, in line with the previously reported observation that peritoneal AAM^res^ have lower proliferation capacity during *Litomosoides sigmodontis* infection in BALB/c mice (10). Furthermore, the expressions of PD-L2 (*Pdcd1lg2*) (Figure 4D) and MHCII molecules (Figure 4E), which are cellular markers typically used to characterize alternative activation (12) in AAM^mono^ of C57BL/6 background, were significantly reduced in BALB/c AAM^mono^. Notably, while 3 of the 5 MHCII genes (*H2-Aa, H2-Ab1, H2-Eb1*) had significantly higher expression in C57BL/6 AAM^mono^, *H2-Ea-ps* expression was specific to BALB/c AAM^mono^. Hence, our studies, together with published findings (10), indicate that macrophages from BALB/c and C57BL/6 mice are functionally distinct in how they respond to IL-4 activation.

### AAMs from C57BL/6 and BALB/c mice respond differently to secondary stimulation by IL-4

Primary stimulation of macrophages could lead to the formation of *de novo* enhancer and transcriptional memory, which consequently hastens the kinetics of activation upon a second repeated stimulation (4, 13). We hypothesized that the chromatin remodeling that occurred with IL-4 stimulation in AAMs may translate into enhanced secondary responses with repeated IL-4 stimulation. To test this hypothesis, we compared the responsiveness of *in vivo* derived F4/80^int^CD206^+^ AAM^mono^ and M^mono^ to a secondary *ex vivo* stimulus of 24 hours (Figure 5A). The secondary stimulation was performed after the harvested macrophages were rested overnight in the absence of IL-4. These differences in secondary responses were also compared between C57BL/6 and BALB/c mice.

**Figure 5:**
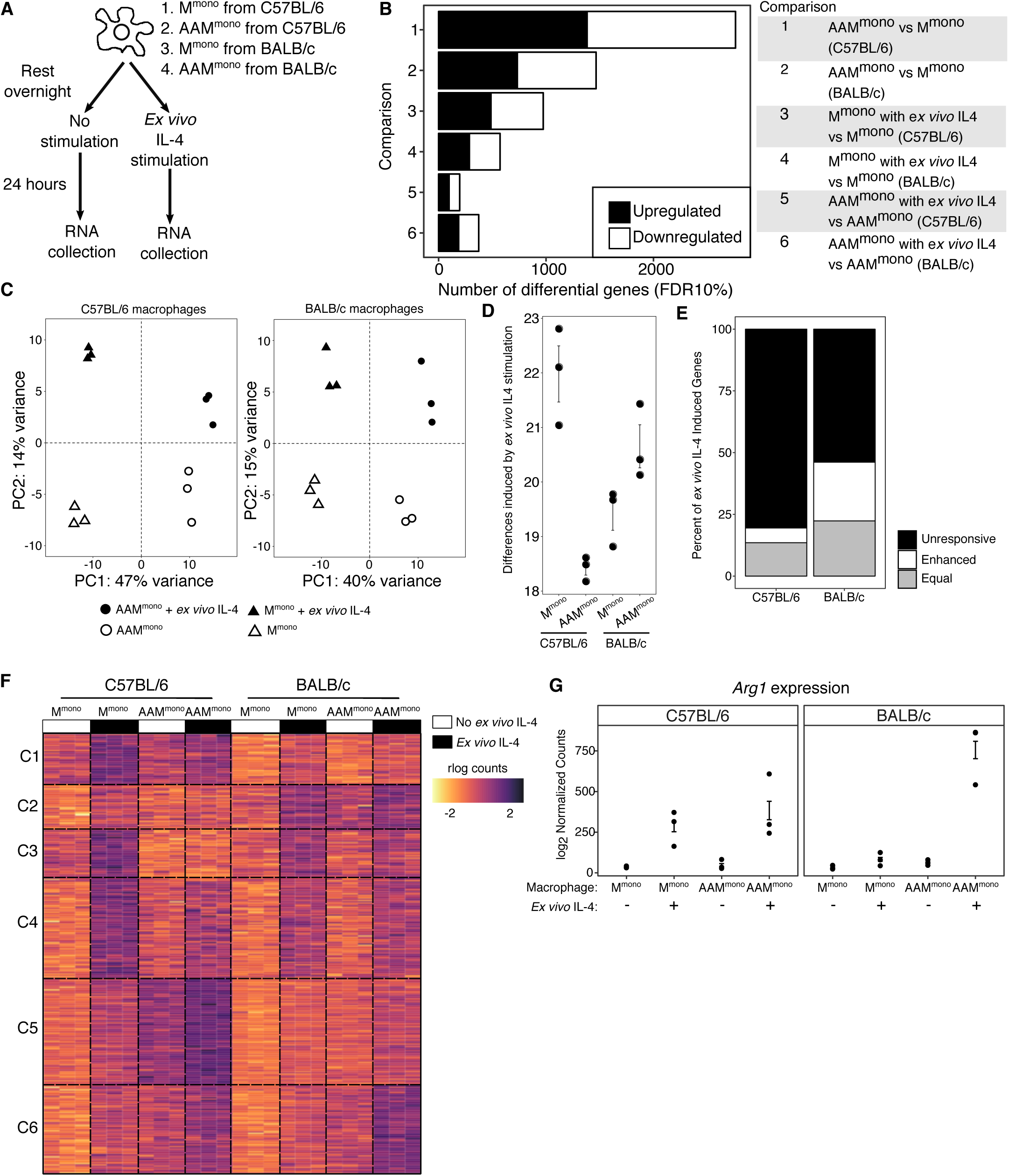
AAMs from C57BL/6 and BALB/c mice respond differently to secondary stimulation by IL-4. (A) Schematic illustrating the study designed to assess differences in response to a secondary IL-4 stimulation between AAMmono and Mmono of C57BL/6 and BALB/c backgrounds. (B) Number of significantly differential genes (FDR < 10%) identified for each comparison. (C) PCA performed separately on 6528 genes with high variance (only genes with variance inter-quartile range > 0.5 were retained) in C57BL/6 and BALB/c macrophages. Data points represent independent biological replicates. (D) Differences in global transcriptional profiles induced by *ex vivo* IL-4, quantitated using Euclidean distance that is calculated between transcriptional profiles (rlog read counts of all non-singleton genes) of cell aliquots from the same cell type that either received a secondary *ex vivo* IL-4 stimulation or cultured in control media. (E) Frequency of genes with enhanced, equal or dampened upregulation after pre-treatment with in vivo IL-4. (F) A union set of 334 genes upregulated by *ex vivo* IL-4 in C57BL/6 and BALB/c macrophages, separated into 6 different clusters by k-means clustering. (G) Expression values of *Arg1* across the different conditions, represented by size-factor normalized read counts. N = 3 mice for each group.

AAM^mono^ from C57BL/6 mice were transcriptionally more distinct from M^mono^ after 24 hours of *ex vivo* culture without IL-4 stimulation, when compared to AAM^mono^ from BALB/c mice (Figure 5B). In general, across both mouse strains, *in vivo* IL-4 stimulation led to greater transcriptional changes than *ex vivo* IL-4 stimulation (Figure 5B-5C). AAM^mono^ were also less responsive to the *ex vivo* IL-4 stimulation, as compared to M^mono^. This indicated that pre-treatment with *in vivo* IL-4 led to reduced response to a secondary *ex vivo* IL-4 stimulation and was contrary to what we had hypothesized. However, the degree of unresponsiveness to the secondary *ex vivo* IL-4 stimulation was more pronounced in AAM^mono^ from C57BL/6 mice than that in BALB/c AAM^mono^ (Figure 5D).

By taking a supervised approach, we determined that 203 of the 252 genes that were inducible by *ex vivo* IL-4 in C57BL/6 macrophages would no longer be responsive if the macrophages were pre-stimulated with *in vivo* IL-4. This was significantly greater (two-sided Fisher’s exact test, p = 1 × 10^−9^) than that observed for BALB/c macrophages, where 111 of the 206 genes that were inducible by *ex vivo* IL-4 would not be responsive after pre-treatment with *in vivo* IL-4 (Figure 5E). Some of the genes that were unresponsive to the secondary IL-4 stimulation were genes that remained persistently upregulated overnight in the absence of IL-4 stimulation (92 of the 203 unresponsive genes in C57BL/6 AAM^mono^ and 39 of the 111 unresponsive genes in BALB/c AAM^mono^). On the other hand, BALB/c macrophages had significantly higher number of genes that were enhanced in upregulation upon IL-4 restimulation (49 of 206 genes in BALB/c vs. 15 of 252 genes in C57BL/6, two-sided Fisher’s exact test, p < 2.2 × 10^−16^) (Figure 5E). When we examined the subset of genes that were only upregulated in BALB/c macrophages with repeated IL-4 stimulation (Figure 5F, Cluster C6), this included *Arg1* (Figure 5F). In AAM^mono^ of C57BL/6 mice, *Arg1* was downregulated overnight in the absence of IL-4 and was equally responsive to *ex vivo* IL-4 stimulation in both M^mono^ and AAM^mono^. In contrast, in macrophages of BALB/c mice, ex vivo *Arg1* expression was only induced with repeated IL-4 stimulations (Figure 5G). Overall, these results support the conclusion that macrophages from BALB/c and C57BL/6 mice are functionally distinct and indicates that this may be particularly important when responding to a secondary stimulation.

## Discussion

In this study, we define and characterize IL-4 induced chromatin accessibility with *in vivo* alternative activation of tissue resident and monocyte-derived peritoneal macrophages. We also show that *in vivo* alternative activation led to a dampening of macrophage response to a repeat stimulation. Hence, *in vivo* IL-4 activation did not only lead to remodeling of the accessible chromatin landscape, but also led to persistent differences in cellular response. However, this dampening of macrophage secondary response occurs to different degrees in mice with different genetic backgrounds.

We propose that PU.1 is one of the key regulators of IL-4 induced chromatin accessibility and PU.1 binding can be mediated through DNA shape readout. DNA shape features, particularly the DNA minor groove width and roll configuration, have been used to distinguish between functional PU.1 binding sites and randomly occurring PU.1 motif (25). However, since DNA shape is a consequence of DNA sequence, both modes of DNA recognition are confounded and difficult to dissociate from one another. For example, it is unclear if the differences of DNA shape in the PU.1 regions of AAM^res^ and AAM^mono^ are simply due to differences in co-factor binding, or if the PU.1 protein in these different lineages of macrophages have different post-translational modifications and recognize binding sites with different DNA shape. Therefore, it is important for future studies to identify which amino acid residue(s) in the PU.1 protein could be involved with DNA shape readout, as mutating these residues could potentially be a strategy to identify PU.1 binding sites that are solely dependent on DNA shape readout, without being confounded by DNA sequences (26).

Several studies have recently described the combinatorial effects of cytokines on macrophage activation, although most of these studies have focused on the opposing effects of type 1 and type 2 cytokines (16, 17). When repeated stimulation with the same cytokine was performed and demonstrated “transcriptional memory” (4, 13), these studies were conducted using short-term stimulation in BMDMs. As such, no comparable studies have so far demonstrated the dampening of response to repeated cytokine stimulation after *in vivo* alternative activation. Whereas an enhanced secondary response in macrophage function is now considered to be a key component of “trained immunity” (27), our results here suggest that a secondary response may also include a dampening of responses under certain circumstances that may be dependent on the genetic background of the host. In future work, it would be important to determine if this dampening of response was specific to *in vivo* IL-4 stimulation, particularly in the setting of a physiological helminth infection model, which should have continuous production of type 2 cytokines at high abundance and also over a chronic infection course. Furthermore, while we have only studied this phenomenon of dampened immune response in AAM^mono^, it should also be investigated in macrophages of other tissue origins, as well as other phagocytes, such as neutrophils and dendritic cells. Finally, given that most human individuals would be exposed to different environmental stimulations throughout their lives, it would be interesting to determine if such repeated stimulations would cause dampening in human macrophage response.

## Materials and Methods

### Experimental methods

#### Mice

Wild type (WT) C57BL/6 mice were purchased from Jackson laboratory and bred onsite for the first set of experiments that compared the effect of IL-4 stimulation on the accessible chromatin profiles from M^res^, AAM^res^, M^mono^ and AAM^mono^. For experiments directly comparing AAMs of C57BL/6 and BALB/c backgrounds, mice of both strains were purchased from Jackson laboratory and immediately used for experiments. Mice were age (7-8 weeks of age) and gender matched. IL-4/anti-IL-4 monoclonal antibody (mAb) complexes (IL-4c) were prepared as described previously (28). To induce AAM^res^, mice were injected intraperitoneally (i.p.) with IL-4c on days 0 and 2. Mice were also treated with 4% thioglycollate alone (to induce M^mono^) or in combination with IL-4c to induce AAM^mono^ (12). All animal procedures were approved by the New York University Institutional Animal Care and Use Committee (IACUC) under protocol numbers 131004 and 130504.

#### Peritoneal cell isolation and cell sorting

Peritoneal cells were isolated by washing the peritoneal cavity twice with cold PBS 1x. Peritoneal exudate were then treated with ACK Lysis buffer to lyse red blood cells and washed once with PBS. Cells were then re-suspended to single-cell suspensions for staining with fluorescently conjugated antibodies at 1:100 dilutions, unless otherwise noted. Antibodies were diluted using 2% fetal bovine serum (FBS). Cells were stained with one of either LIVE/DEAD™ Blue (Invitrogen) or LIVE/DEAD™ Near-IR (Invitrogen), blocked with 4µg/ml anti-CD16/32 (2.4G2; Bioxcell) and stained with anti-CD11b Pacific Blue (M1/70; Biolegend), F4/80 PECy7 (BM8; Biolegend), CD206 APC (C068C2; Biolegend), Siglec-F PE (E50-2440; BD Biosciences), CD3 PE (145-2C11; Biolegend), CD19 PE (6D5; Biolegend), CD49b PE (DX5; Biolegend), Ly6G (1A8; Biolegend), PD-L2 (PerCP-Cy55; Miltenyi; diluted at 1:20), MHCII (APC-Cy7; Biolegend). Cells were gated on singlet, live, Dump-negative (CD3-, CD19-, DX5-, Siglec-F-, Ly6G-), CD11b+, then subsequently gated on their M^res^ and AAM^res^ (F4/80hi, CD206-) or M^mono^ and AAM^mono^ (F4/80int, CD206+) phenotype. Cell surface expression of PD-L2 and MHCII were acquired for analysis. Cells were sorted using 100µm nozzle into FBS, on either BD FACSAriaII or SONY HAPS1, depending on instrument availability.

#### Assay for Transposase-Accessible Chromatin with Sequencing (ATAC-seq)

ATAC-seq was performed as described by Buenrostro et al (18). 50,000 FACS-purified cells per sample were spun down at 400g for 5 min at 4°C and washed once with 50µl cold PBS. Cells were lysed with 50µl lysis buffer (10 mM Tris-HCl, pH7.4, 10 mM NaCl, 3 mM MgCl2, 0.1% IGEPAL CA-630) and immediately spun down at 1500rpm for 10 min at 4°C. The isolated cell nuclei were then incubated for 30 min at 37°C with 50µl of transposase reaction, which contained 25µl Tagment DNA buffer (Illumina), 2.5µl Tagment DNA enzyme (Illumina) and 22.5µl nuclease-free water. The transposed DNA was immediately purified using the Qiagen MinElute PCR Purification Kit (Qiagen) following manufacturer’s guide and eluted at 10ul volume. PCR amplification of the transposed DNA was done using a low-cycle number protocol and with primers published by Buenrostro et al (18). Each PCR mix contained of 25µl of NEB 2x PCR Mix (New England Biolabs), 2.5µl of 25µM forward primer (Primer Ad1_noMX), 2.5µl of 25µM reverse barcoded primer, 0.3µl of 100x SYBR Green (Invitrogen) and 10µl of transposed DNA. PCR was carried out with the cycling protocol: 72°C for 5 min, 98°C for 30s, followed by 5 cycles of 98°C for 10s, 63°C for 30s, 72°C for 1 min. The reaction was held at 4°C after the 5^th^ cycle. A side qPCR was set up using the PCR product from these 5 cycles of amplification. Each qPCR mix contained 5µl NEB 2x PCR Mix, 0.25µl 25µM forward primer, 0.25µl 25µM reverse barcoded primer, 0.06µl 100x SYBR Green, 4.44µl nuclease-free water and 5µl of the PCR-amplified product. qPCR was carried out using the cycling protocol: 98°C for 30s, followed by 25 cycles of 98°C for 10s, 63°C for 30s, 72°C for 1 min and plate read. The qPCR amplification plot was then used to calculate the additional number of cycles needed for the PCR to achieve maximum amount of product without going into saturation. Each sample was amplified for a total of 14-16 cycles. The amplified libraries were then purified using Qiagen MinElute PCR Purification kit following manufacturer’s guide and eluted at 20µl volume. Libraries were sequenced on the HiSeq 2000 with 2 x 50 cycles and for an average of 50 million paired-end reads per sample. We performed the IL-4c stimulation experiment twice and generated two independent sets of libraries to obtain an optimal number of biological replicates for each macrophage population. The two independent sets of libraries are referred here after as “Run 1” and “Run 2”, respectively. ATAC-seq libraries for C57BL/6 and BALB/c AAMs were generated using the same protocol and sequenced in a single run.

#### Transcriptional profiling of BALB/c and C57BL/6 AAMs

100,000 cells were sorted per sample as described above. FACS-purified cells were spun down and washed once with PBS before lysis with 350µl of Buffer RLT from the RNeasy Mini Kit (QIAgen). RLT lysates were homogenized by 1 minute of vortexing and were immediately stored at −80°C until RNA isolation. RNA was isolated using the RNeasy Mini Kit (QIAgen) based on manufacturer’s protocol, with an additional DNase digestion step using the RNase-free DNase set (QIAgen). Transcriptional profiling was done using the CEL-seq2 protocol (29) and library preparation was performed at the NYU School of Medicine Genome Technology Center core facility. CEL-seq2 libraries were sequenced on the HiSeq 4000 with 2 x 50 cycles. While CEL-seq2 was originally developed as a single-cell assay, we used this protocol in this study as a bulk transcriptional profiling assay and use the more commonly-used terminology “RNA-seq” to describe data generated from this assay.

#### Re-stimulation experiment

300,000 cells (M^mono^ or AAM^mono^ from C57BL/6 and BALB/c mice, induced using the injection protocols described above) were sorted per sample as described above. FACS-purified cells were washed twice, then re-suspended to 1ml PBS for cell counting on the hemocytometer. Cells were spun down again and re-suspended to a single-cell suspension of 100,000 cells per 200µl, using DMEM with 10% FBS and 1% Penicillin/Streptomycin. Two aliquots of 100,000 cells were obtained from each sample and plated separately onto wells of a 48-well tissue-culture plate in 200µl. Cells were incubated overnight (approximately 18 hours) at 37°C, 5% CO_2_. After the overnight incubation, the initial culture media was discarded and replaced with fresh media. An *ex vivo* dose of IL-4 (20ng/ml) was added for designated wells. Control wells received fresh media only. After 24 hours of stimulation, media was aspirated completely from each well and 350µl of Buffer RLT was immediately added for cell lysis. The lysed cells of each sample was transferred to a 1.5ml Eppendorf tube, vortexed for 1 minute and immediately stored at −80°C until RNA isolation. RNA isolation and transcriptional profiling by CEL-seq2 were performed as described above.

### Bioinformatics and computational methods

#### ATAC-seq sequence processing

Raw ATAC-Seq reads were aligned to the reference mouse genome mm9 using bowtie2 (v2.2.9) (30), with the parameters --maxin 2000 and --local, while keeping all other parameters at default settings. To keep only highly unique alignments, reads with MAPQ score less than 30 were removed. We further removed all duplicate reads, as well as reads mapping to mitochondrial DNA and chromosome Y. Read filtering steps were done using the suite of tools from samtools (v1.2 and v1.3.1) (31), ngsutils (v0.5.9) (32) and picard-tools (http://broadinstitute.github.io/picard/, v1.1.1 and v2.8.2). After all filtering steps, reads were merged across all replicates from the same macrophage population. This resulted in a median depth of 15,235,324 reads per macrophage population in Run 1 and 9,865,310 reads per macrophage population in Run 2. For visualization of accessibility reads on the Integrative Genomics Viewer (IGV), we merged reads from the same macrophage population across samples from both runs, generated tiled data format (TDF) files using IGVtools and finally normalized the merged reads to reads per million (RPM) (33).

#### Identification of accessible chromatin regions

We used the merged reads for each macrophage population to identify accessible chromatin regions, using the PeaKDEck (v1.1) peak calling algorithm, which measures signal density from randomly sampled bins genome-wide before generating a data set-specific probability distribution to identify regions with significant signal enrichment (34). We ran PeaKDEck using sampling bins that consist of a 75bp central bin (-bin) and a 10000bp background bin (-back). Sampling along the genome was done in steps (-STEP) of 25bp and the background probability distribution was generated using 100000 randomly selected sites (-npBack). Significance was defined using a p-value of less than 0.0001 and regions with significant p-values were defined as a “peak” (i.e. an accessible chromatin region). Peak calling was done independently on libraries generated from Run 1 and Run 2.

#### Generation of a union set of accessible chromatin regions

We next counted the number of reads present at each accessible region in order to analyze the ATAC-Seq data using quantitative approaches downstream. To do this, we first generated a set of consensus peaks across the data set by taking the union of peaks called from each macrophage population. Peaks were merged if overlapping by 1bp or more. The number of reads at each peak within the union peak sets were then counted for each sample. Finally, each peak was re-centered ±100bp on its summit, defined as the position with maximum pile up of reads. Re-centering on peak summits was performed as this should coincide with the binding event of a transcription factor within an accessible chromatin region. We implemented the read counting and peak summit re-centering steps directly using the dba.count function from the Bioconductor package DiffBind (version 1.14.2) (35). The final count matrix, which consisted of 61,713 peaks, was used for downstream analyses.

#### Quantitative ATAC-seq analysis

ATAC-seq read counts were transformed using the regularized logarithmic (rlog) transformation as implemented in the Bioconductor package DESeq2 (36). To manage batch effect from the two separate libraries, we first modeled the rlog accessibility read counts to the batch variable using a linear model and subtracted out the coefficient contributed by the batch variable – this was implemented directly using the removeBatchEffect function in limma (37). We next chose a set of 30,856 regions with high variance, using the varFilter function in the genefilter package with default parameters, which keeps only features with variance inter-quartile range > 0.5 (38). We performed principal component analysis (PCA) using the batch-subtracted rlog read counts of these regions with high variance using the prcomp function in R.

To identify IL-4 dependent accessible regions, we directly compared the ATAC-seq profiles of IL-4 stimulated macrophages to their reference non-stimulated macrophages, using a differential analyses workflow directly implemented through DESeq2. We fit the negative binomial model in DESeq2 using the raw accessibility reads from all 61,713 regions, with the model **∼** Batch + Population, where Batch is a variable describing if a sample belonged in Run 1 or Run 2, while Population is a variable describing if the sample is M^res^, AAM^res^, M^mono^ or AAM^mono^. IL-4 dependency was defined using a significance threshold of False Discovery Rate (FDR) of 10%. To visualize IL-4 dependent regions, we scaled the batch-subtracted rlog read counts of these IL-4 dependent regions by z-score transformation and next performed k-means clustering on these scaled, rlog-transformed reads (K = 4). The clustered matrix was visualized as a heatmap.

#### Comparison of sequence properties between constitutively accessible and IL-4 induced regions

##### Identification of constitutively accessible and IL-4 induced regions

To define a set of constitutively accessible regions, we used only peaks from M^res^ and M^mono^, respectively, that were identified in both Run 1 and Run 2. This resulted in 8061 constitutively accessible regions in M^res^ and 14,045 constitutively accessible regions in M^mono^. IL-4 induced peaks were defined using the differential analysis outlined above. All region overlap analyses throughout this study were performed using the intersect function from the BEDTools suite (39) and overlaps were defined as any regions overlapping by at least 1bp, unless otherwise noted.

##### Genomic elements enrichment analysis

We downloaded genome-wide annotations of five different genomic elements (promoter, start exon, coding exon, end exon, intron) from the UCSC Known Gene database for mm9 (40). We defined promoter elements as the 200bp-region upstream of a transcriptional start site (TSS). We next assigned each of the 61,713 accessible regions in our data set to a unique genomic element label. Where an accessible region overlapped two different types of genomic elements, we assigned it to the element with higher number of overlapping base pairs. Finally, any chromatin regions not assigned to one of these five genomic elements were labeled as intergenic. To determine the enrichment levels of a particular type of genomic element *G* within a given set of accessible regions *A* (either constitutively accessible or IL-4 induced regions in AAM^res^ or AAM^mono^), we used the binomial cumulative probability distribution, b(*x*; *n,p*), where *x* = number of success, *n* = number of trials and *p* = background probability of success. We used the pbinom function in R. We defined *x* to be the number of accessible regions in *A* that were labeled as the genomic element *G* that was being tested, *n* to be the total number of genomic elements *G* detected in our combined data set and *p* to be the proportion of the accessible region *A* to the total 61,713 accessible regions. This then gave the enrichment levels of *G* in *A*, relative to all the accessible regions identified across the different macrophage populations.

##### G/C content analysis

To calculate percentage GC, we first used the hgGcPercent function from the kentTools suite (v20170111, UCSC Genome Bioinformatics Group, https://github.com/ucscGenomeBrowser/kent) to quantitate the number of G or C bases in each accessible region. This value was next normalized using the length of the accessible region. CpG island track was downloaded from the UCSC Genome Annotation Database for mm9 (http://hgdownload.soe.ucsc.edu/goldenPath/mm9/database/). Enrichment levels of CpG island in a given set of accessible regions *A* was based on the binomial cumulative probability as described above, where *x* = number of accessible regions in *A* that overlapped a CpG island, *n* = number of CpG island in the total data set of 61,713 regions and *p* = proportion of *A* to the total 61,713 regions.

##### Calculation of distance to IL-4 induced genes

IL-4 induced genes were first identified for AAM^res^ and AAM^mono^ from the microarrays generated by Gundra and Girgis *et al* (12), using the linear model and empirical Bayes statistics as implemented in limma, with genes significantly induced by IL-4 defined using the thresholds FDR 10% and log_2_ fold change > 1. The distance between each accessible region and its closest IL-4 induced gene body was calculated using the closest function in BEDTools.

#### Transcription factor (TF) motif analysis

Sequences of IL-4 induced regions were fetched using the BEDTools getfasta function for TF motif analysis with the MEME Suite tools (41). Whole genome fasta file for mm9 was downloaded from the Illumina igenome database (https://support.illumina.com/sequencing/sequencing_software/igenome.html). Background file was generated using the function fasta-get-markov in MEME, based on the total 61,713 accessible regions at a Markov model order of 3. TF motif databases (which included mouse and human TF motifs) were curated as described in (42). We performed *de novo* motif discovery by running MEME (as part of MEMEChIP, which randomly sampled 600 sequences) with the parameters: -mod zoops -nmotifs 3 -minw 6 -maxw 30. Over-representation analysis to identify macrophage-specific TF motifs was performed by running HOMER (43) using sequences from the opposing macrophages as background sequences (i.e. to identify TF motifs specific to AAM^res^, sequences of IL-4 induced regions from AAM^mono^ were used as background sequences) and the parameters: -mask, -size 8,10,12,16, -mset vertebrates, -nlen 3. For motifs from each macrophage lineage, we combined all three *de novo* motifs discovered by MEME and motifs with enrichment log_2_ p-values < −15 by HOMER for clustering analysis (we used the known motifs output from HOMER). This resulted in 14 motifs for AAM^res^ and AAM^mono^, respectively. Clustering was done using STAMP (44, 45), with the frequency matrices of motifs and the default parameters of: column comparison metric – Pearson correlation coefficient, alignment method – ungapped Smith-Waterman, tree-building algorithm – UPGMA, multiple alignment strategy – iterative refinement.

#### Comparisons between predicted PU.1 motif and ChIP-seq defined PU.1 binding sites

PU.1 ChIP-seq regions identified in thioglycollate-induced macrophages, generated from two different experiments, were downloaded as BED files that had been directly deposited on Gene Expression Omnibus (GEO) (GSM1131238 and GSM1183968) (5, 21). A set of 55,386 reproducible PU.1 binding sites were defined by intersecting these two sets of PU.1 ChIP-seq regions. We used the PU.1 motif discovered *de novo* from all the constitutively accessible regions in M^mono^ and ran FIMO to identify all PU.1 motif sites from M^mono^, using a p-value threshold of 0.0001 and the background file generated as described above. Since the published PU.1 ChIP-seq regions were of 200bp length, we extended the predicted PU.1 motifs from M^mono^ by ±100bp to match the comparison. Overlapping rate was calculated as total number of predicted PU.1 motifs from M^mono^ overlapping a reproducible PU.1 ChIP-seq region / total number of predicted PU.1 motifs from M^mono^ × 100%.

#### PU.1 motif analysis

We identified IL-4 induced PU.1 motif sites by performing motif scanning with FIMO (46), using the PU.1 motif discovered *de novo* from the IL-4 induced peaks of AAM^res^ and AAM^mono^, respectively. FIMO was run with a p-value threshold of 0.0001 (as part of MEMEChIP). Motif scores were also calculated as part of FIMO.

The detected IL-4 induced PU.1 motif sites were then extended ±25bp using the BEDTools slop function. These PU.1 motif sites ± 25bp are hereafter referred to as “PU.1 regions”. To identify potential co-factors for PU.1, these PU.1 regions were specifically subjected to motif scanning by FIMO, with a background model that was based on the 55,386 PU.1 ChIP-seq peaks described above and at a Markov model order of 3. To determine which of these detected motifs were macrophage-specific, the odds ratio of a motif being detected in the PU.1 regions of AAM^res^ vs. AAM^mono^ were calculated. P-value was determined using two-sided Fisher’s test, with the null hypothesis of a motif being equally likely to be detected in the IL-4 induced PU.1 regions of AAM^res^ and AAM^mono^ (i.e. log_2_ odds ratio of zero). When multiple motifs of the same TF were present in the database, we used the motif that was most frequently detected in PU.1 regions for odds ratio calculations. TFs from mouse and human were kept as separate analyses, but included in the same visualization. To account for the multiple hypotheses testing performed over 633 mouse TFs and 835 human TFs, the Benjamini-Hochberg procedure was used to perform p-value adjustment by calculating the FDR and significance threshold was set at FDR 10%. Hence, statistically significant TFs were macrophage-specific, with log_2_ odds ratio > 0 indicating AAM^res^-specificity and log_2_ odds ratio < 0 indicating AAM^mono^-specificity. For visualization of significant results, TFs were summarized at the family level as defined in (47) (Figure 3B). The maximum absolute log_2_ odds ratio of the family was visualized and the specific TF with the maximum absolute log_2_ odds ratio value was stated in parenthesis. Where TF family annotation was not available, the log_2_ odds ratio of the specific TF itself was used.

DNA shape features of PU.1 motifs were analyzed using the DNAshape algorithm (24, 48) for 4 different DNA shape configurations at single nucleotide resolution. Sequences on the anti-sense strand were reverse complemented prior to DNA shape prediction.

#### Processing of CEL-seq reads

CEL-seq reads were first demultiplexed using the bc_demultiplex script from https://github.com/yanailab/CEL-Seq-pipeline (29). Demultiplexed reads were aligned to the mm10 mouse reference genome using bowtie2 (version 2.2.9). Aligned reads were counted for each gene using a modified htseq-count script (from https://github.com/yanailab/CEL-Seq-pipeline) adapted for CEL-seq reads with unique molecular identified (UMI). We included only reads with MAPQ score > 30 and removed singleton genes. This resulted in a final median read depth of 737,848 reads per sample, covering a median of 11,096 genes per sample. PCA was performed using 7431 genes with high variance, defined using the varFilter function in the genefilter package with default parameters, which keeps only features with variance inter-quartile range > 0.5. Differential analysis was done using DESeq2 by fitting the negative binomial model using ∼ Strain + CellType + Strain:CellType. Significantly differential genes were extracted using a threshold of FDR 10% for the four different comparisons of: (1) AAM^res^ vs. AAM^mono^ in C57BL/6, (2) AAM^res^ vs. AAM^mono^ in BALB/c, (3) BALB/c vs. C57BL/6 in AAM^res^ and (4) BALB/c vs. C57BL/6 in AAM^mono^. Overlapping genes were defined as genes identified as differential in two different comparisons. Pathway enrichment analysis was done using strain-specific genes in AAM^res^ and AAM^mono^, respectively, through Ingenuity Pathway Analysis (IPA) with the parameter Organism = Mouse and keeping all other parameters at default settings.

#### Defining gene classes from re-stimulation experiment

CEL-seq reads were processed as described above. This resulted in a final median read depth of 979,547 reads per sample, covering a median of 10,820 genes per sample. PCA was separately performed on the rlog read counts from 6528 high variance genes from C57BL/6 and BALB/c, respectively. The differences in transcriptional profiles induced by *ex vivo* IL-4 stimulation were represented as Euclidean distances, which were calculated between transcriptional profiles (rlog read counts of all non-singleton genes) of cell aliquots from the same cell type that either received a secondary *ex vivo* IL-4 stimulation or cultured in control media. Differential analysis was done using DESeq2 by fitting the negative binomial model using ∼ Group, where Group is a factor consisting of 8 different levels (M^mono^ from C57BL/6, AAM^mono^ from C57BL/6, M^mono^ with *ex vivo* IL-4 stimulation from C57BL/6, AAM^mono^ with *ex vivo* IL-4 stimulation from C57BL/6, M^mono^ from BALB/c, AAM^mono^ from BALB/c, M^mono^ with *ex vivo* IL-4 stimulation from BALB/c, AAM^mono^ with *ex vivo* IL-4 stimulation from BALB/c). Significantly differential genes were extracted using a threshold of FDR 10% and the comparisons are described on Figure 7B.

To define “Unresponsive”, “Enhanced” and “Equal” genes, we identified genes that were either uniquely or commonly differential in the comparisons “AAM^mono^ with *ex vivo* IL-4 vs. AAM^mono^” and “M^mono^ with *ex vivo* IL-4 vs. M^mono^” (49). “Unresponsive” genes were defined as genes that were uniquely differential in “M^mono^ with *ex vivo* IL-4 vs. M^mono^”, while “Enhanced” genes were uniquely differential in “AAM^mono^ with *ex vivo* IL-4 vs. AAM^mono^”. For genes that were differential in both comparisons, we further examined the magnitude of fold change – if the fold change (defined as a differences in log_2_ fold change of >1) is greater in the “M^mono^ with *ex vivo* IL-4 vs. M^mono^” comparison, it is defined as an “Unresponsive” genes and if the fold change is greater in the “AAM^mono^ with *ex vivo* IL-4 vs. AAM^mono^” comparison, it is defined as an “Enhanced” gene. Finally, if the magnitude of fold change is similar in both comparisons (defined as a differences in log_2_ fold change of <1), the gene is defined as “Equal”. This categorization process of different gene classes was performed for C57BL/6 and BALB/c macrophages, respectively. For visualization of these different gene groups, the rlog read counts from all of these genes were extracted, scaled to a mean of 0 and standard deviation of 1 across each sample, and subjected to k-means clustering, with K=6. Heatmap was visualized using the R package pheatmap.

## Supporting information

DataS3

## Acknowledgements

We thank the NYU School of Medicine Genome Technology Center and Cytometry and Cell Sorting Laboratory core facilities. These shared resources are partially supported by the Laura and Isaac Perlmutter Cancer Center support grant P30CA016087. We also thank the NYU IT High Performance Computing resources, services, and staff expertise. This work was supported through the NIH, NIAID grants AI093811 and AI094166 (P.L.) and NIDDK grant DK103788 (P.L.), American Association of Immunologists (M.S.T.), Vilcek Foundation (M.S.T.).

## Author contributions

Conceptualization, M.S.T. and P.L.; Methodology, M.S.T, E.R.M., N.M.G., R.A.B. and P.L.; Formal Analysis, M.S.T.; Investigation, M.S.T. and N.M.G.; Writing – Original Draft, M.S.T. and P.L.; Writing – Review and Editing, E.R.M. and P.L.; Visualization, M.S.T.; Data Curation, M.S.T.; Supervision, R.A.B. and P.L.; Funding acquisition, P.L. All authors read and approved of the final draft.

## Data availability

ATAC-Seq and RNA-Seq data have been deposited on NCBI database Gene Expression Omnibus (GEO) under the SuperSeries GSE116108 (ATAC-Seq subseries GSE116107 and RNA-Seq subseries GSE116105).

## Figures legends

**Supplemental Figure 1:**
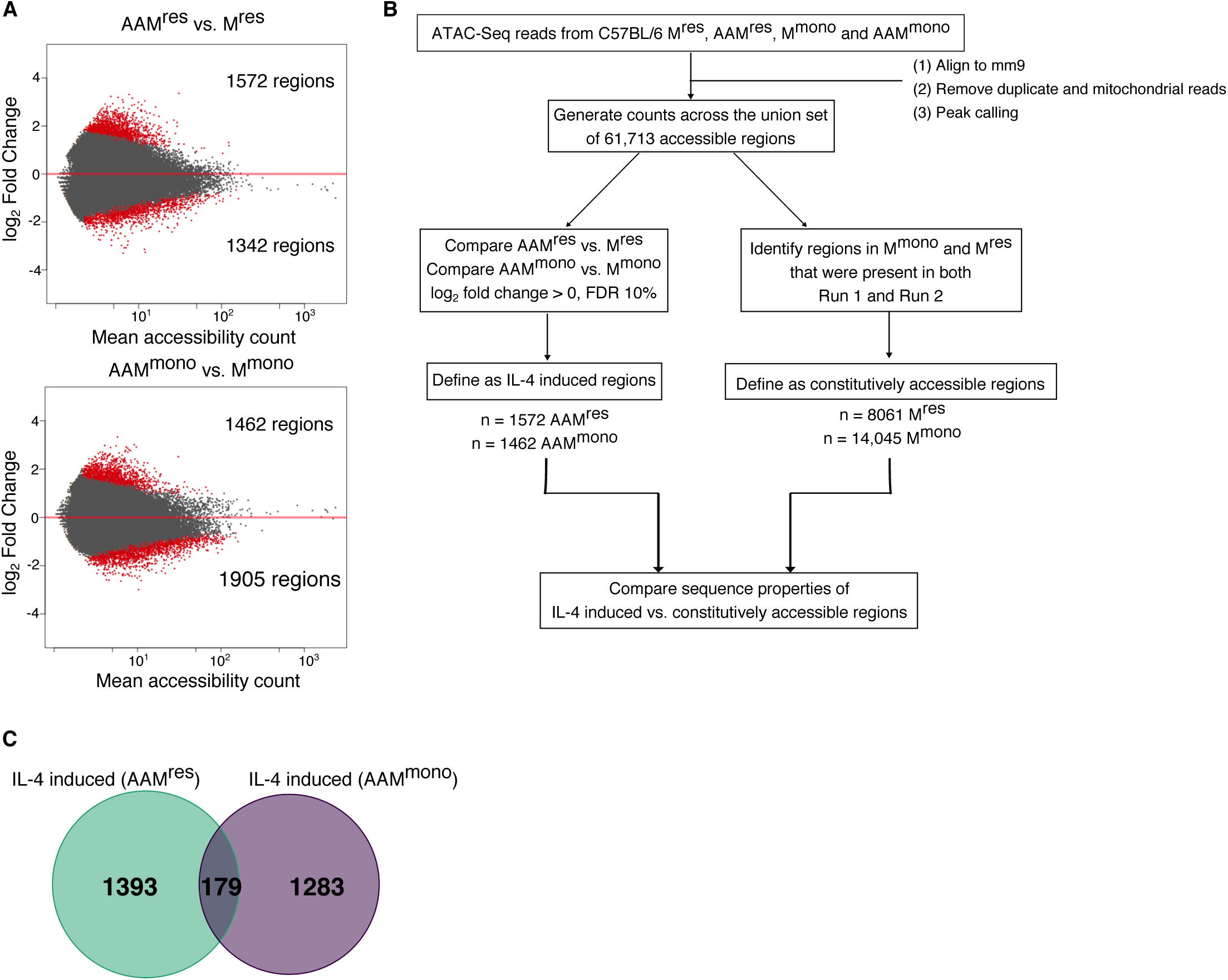
IL-4 stimulation leads to remodeling of open chromatin landscape in peritoneal macrophages. (A) Comparisons between the accessible chromatin profiles of IL-4 stimulated macrophages and non-stimulated macrophages, presented as MA plots (left – AAM^res^; right – AAM^mono^). Differential chromatin regions (FDR 10%, |LFC| > 0) are highlighted in red. (B) Venn diagram illustrating the minimal overlap between IL-4 induced regions in AAM^res^ and AAM^mono^. Values in Venn diagrams represent the number of unique accessible regions in each corresponding set. (C) Schematic outlining the workflow to identify IL-4 induced regions and constitutively regions for comparisons of the sequence properties associated with these two classes of genomic elements.

**Supplemental Figure 2:**
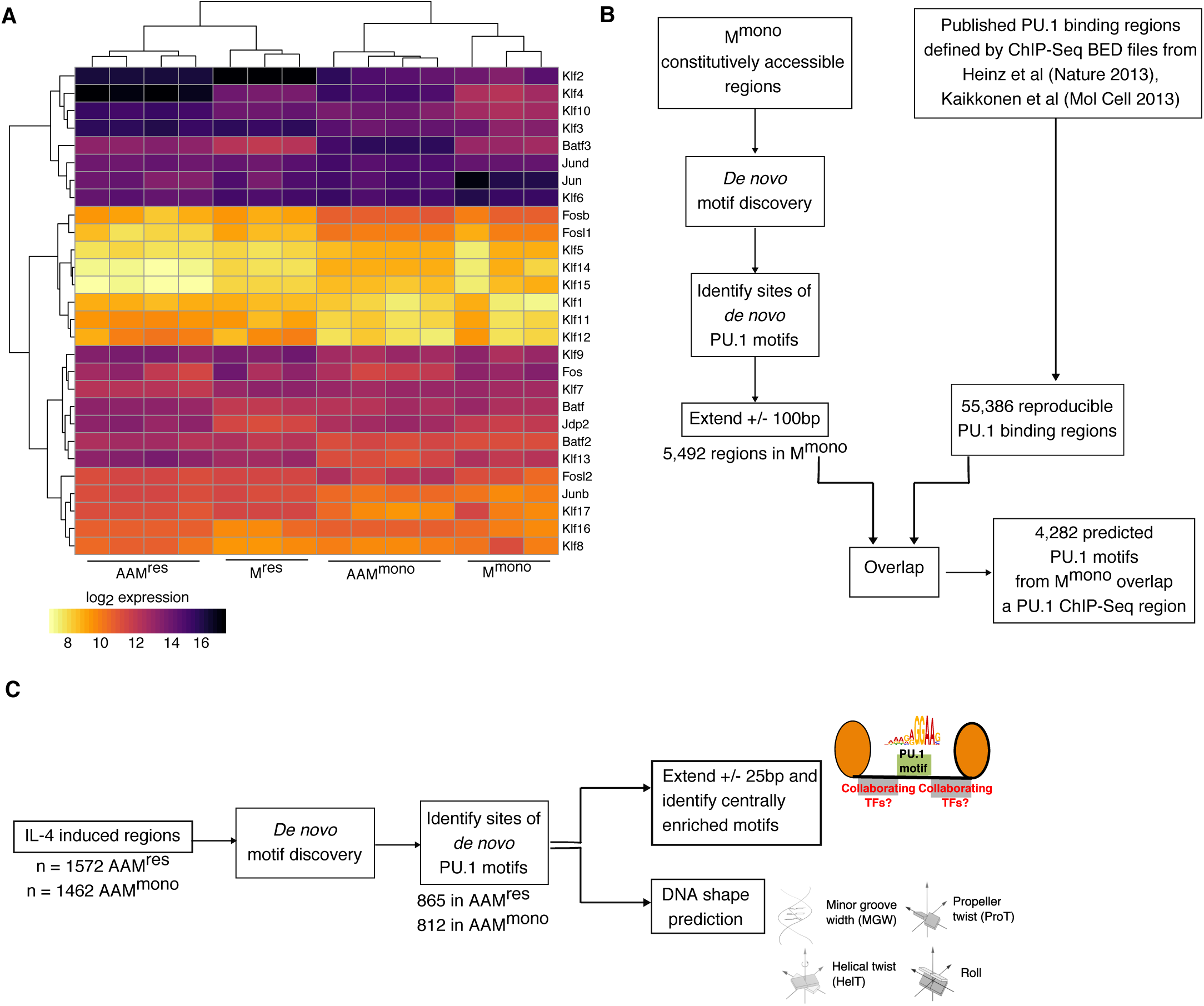
Motif analyses of IL-4 induced regions in AAM^res^ and AAM^mono^. (A) Expression values of all TF genes related to the *de novo* motifs discovered from the IL-4 induced regions. Values are log_2_ intensity values of microarrays (12) and were clustered by hierarchical clustering. TF genes falling into the 2^nd^ cluster were excluded from the analysis presented in Figure 2D due to their low expression. (B) Schematic outlining the workflow used to verify the accuracy of predicted PU.1 motif sites by comparing to published ChIP-Seq data sets. (C) Schematic outlining the workflow used to analyze features of IL-4 induced PU.1 motif sites.

**Data S1.** Complete lists of TF motifs and associated statistics identified by the over-representation approach. Related to Figure 2E.

**Data S2.** Complete list of macrophage-specific motifs detected within ±25bp of PU.1 motifs in AAM^res^ and AAM^mono^. Related to Figure 3B.

**Data S3.** Composite plots of predicted DNA shape values at single nucleotide resolution, between base pair 3-13 of the PU.1 motifs in AAM^res^ and AAM^mono^. Related to Figure 3D.

**Data S4.** List of pathways enriched among strain-specific genes from AAM^res^ and AAM^mono^. Related to Figure 4C.

