## Supplementary figures and images for "Alternative activation of macrophages is accompanied by chromatin remodeling and short-term dampening of macrophage secondary response"

### DataS3

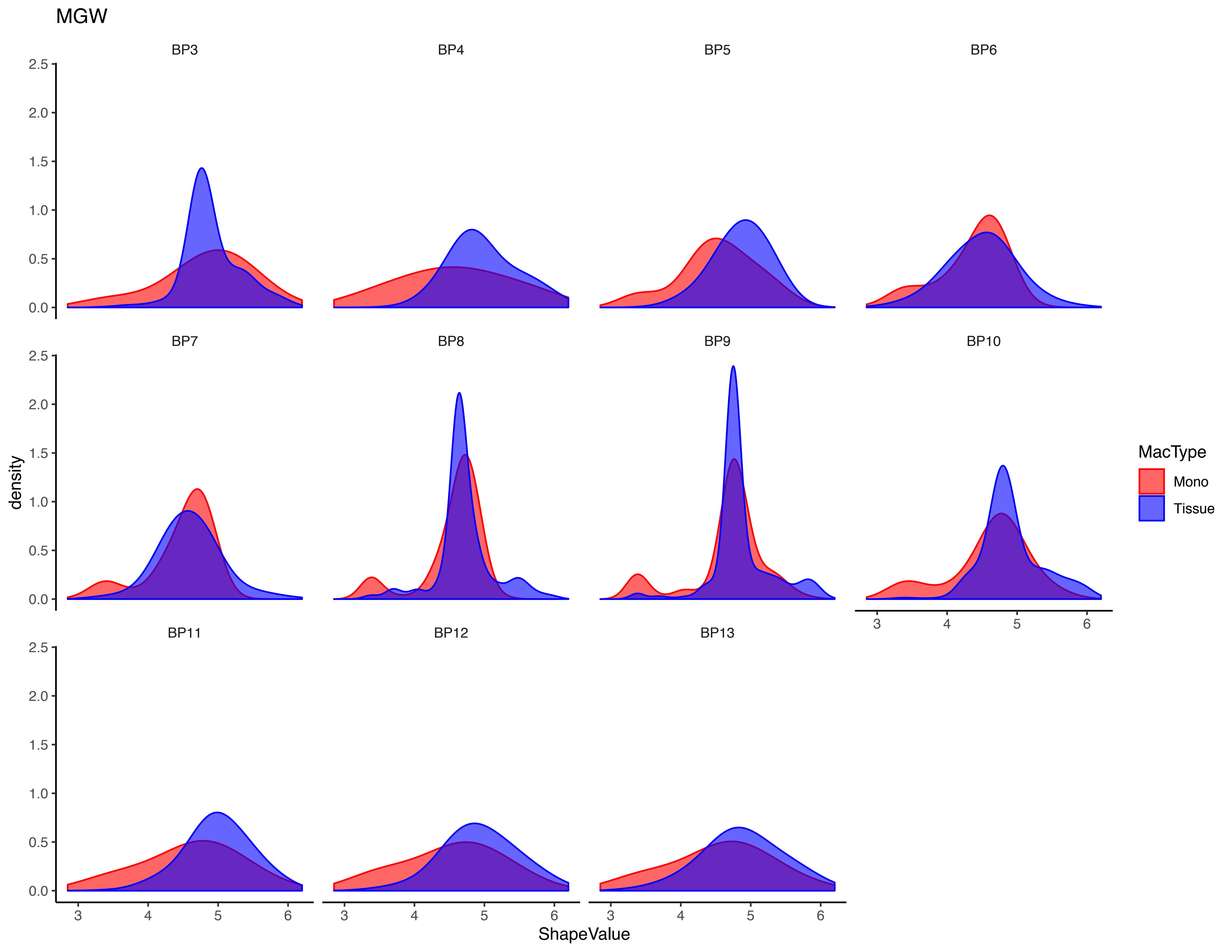

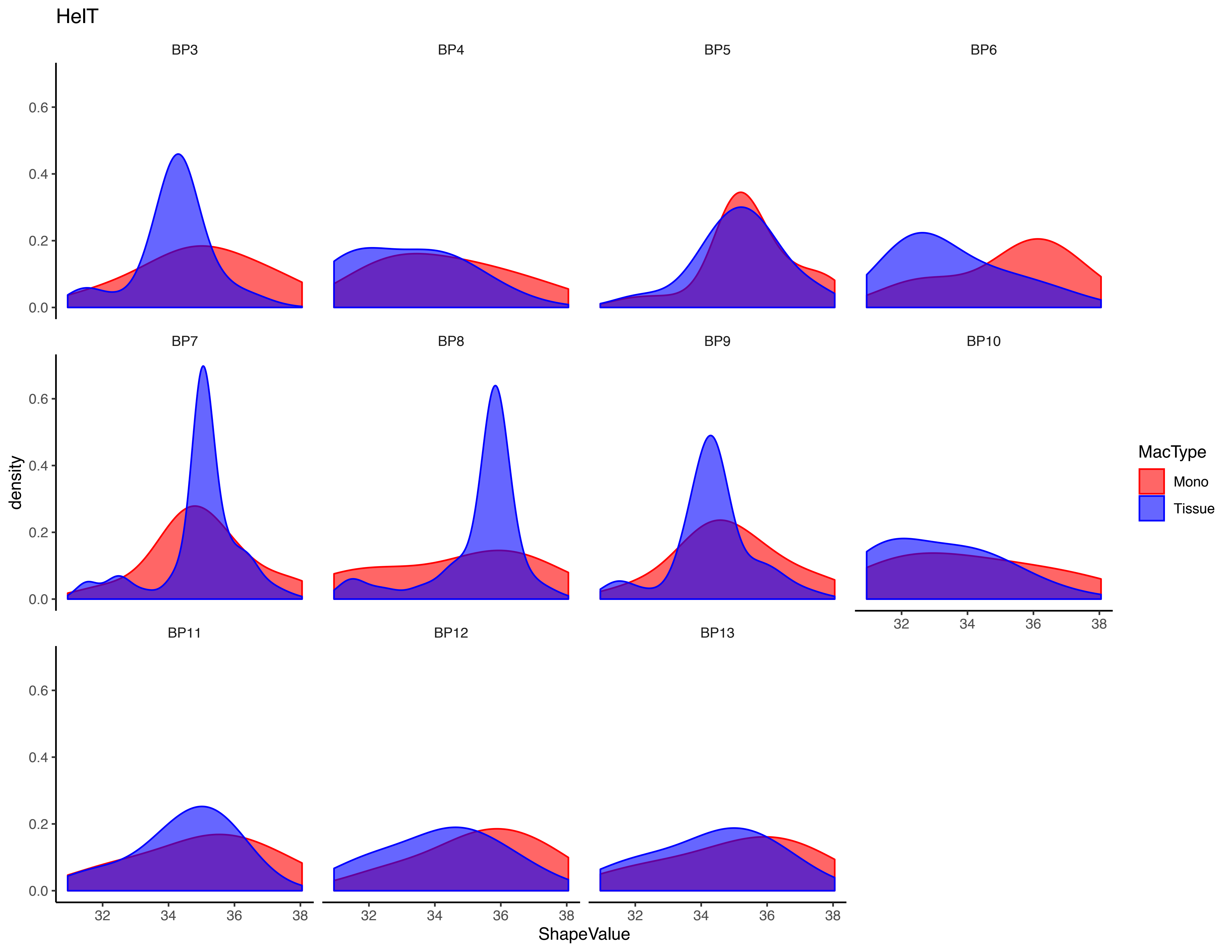

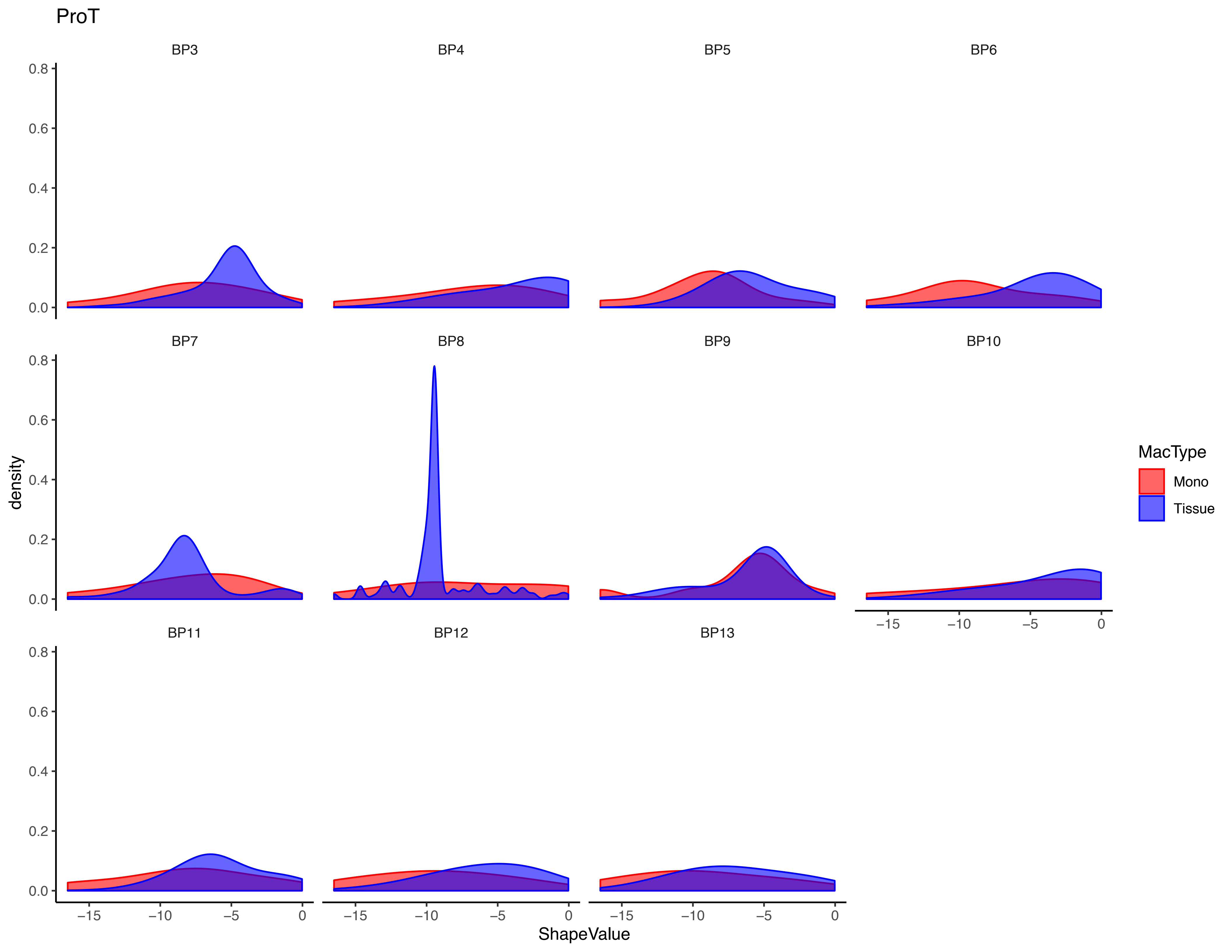

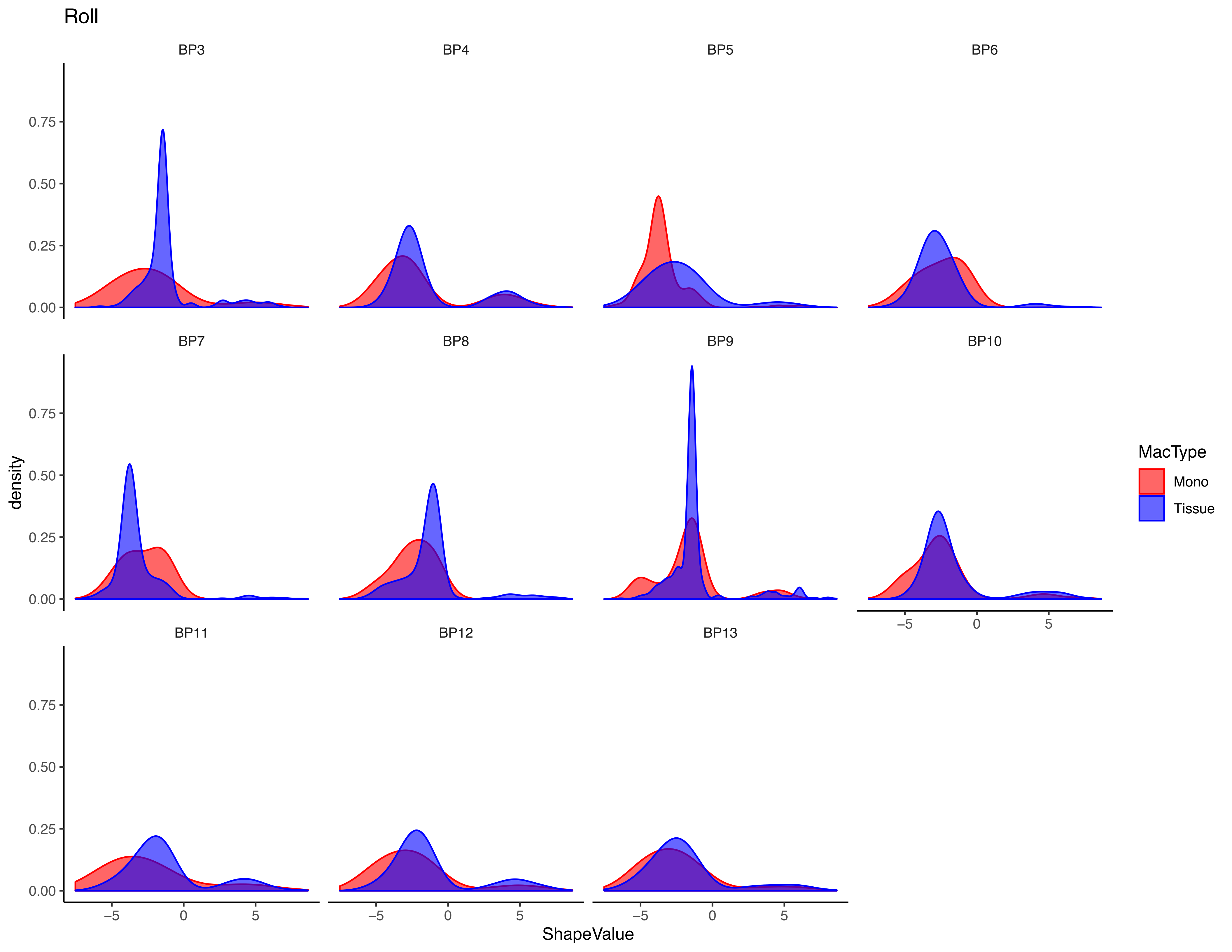
